# Postpartum enhancement of spatial learning and cognitive flexibility: an IntelliCage study

**DOI:** 10.1101/2025.09.27.678974

**Authors:** Melinda Cservenák, Luca Darai, Tamara C. Kállai, Bori Záhonyi, Norbert Bencsik, László Détári, Arpád Dobolyi

**Author notes:** Corresponding author: Arpád Dobolyi, PhD, Department of Physiology and Neurobiology, Eötvös Loránd University, Budapest, Hungary /8775.

## Abstract

The transition to motherhood has been shown to result in significant changes in the structure and function of the brain, with particular emphasis on the enhancement of cognitive abilities that are essential for survival. Specifically, in murine models, spatial learning and cognitive flexibility have been identified as critical components of mothering. These abilities facilitate efficient navigation of the environment, resource acquisition, and responsiveness to offspring’s needs. While cognitive enhancements during the postpartum period have been observed in various experimental setups, research using long-term data collection with automated monitoring in home cage setup is completely lacking even though it provides a more reliable approach than other experimental procedures. This study aims to address this knowledge gap by systematically examining spatial learning and cognitive flexibility in female mice during the reproductive stages using IntelliCage, with the objective of offering a more comprehensive understanding of maternal cognitive adaptations. Utilising the IntelliCage system, we observed that female mice in the postpartum phase outperformed pregnant and nulliparous females in place learning, reversal learning, and fixed schedule drinking tasks, demonstrating faster adaptation and superior retention of information. These paradigms mirror real-world challenges faced by mothers, such as navigating resources and balancing caregiving with self-maintenance. The enhanced performance demonstrated by the mothers could be attributed to their heightened motivation and cognitive abilities, potentially influenced by substantial hormonal shifts, which have been known to modify neuroplasticity in critical brain regions. The identification of these improvements in maternal behaviour may offer novel insights into the impact of reproductive experiences on brain function, with implications for maternal health and broader cognitive research.

## Introduction

The postpartum period is characterized by profound physiological, emotional, and behavioral transformations aimed at optimizing maternal care for offspring survival (Agrati & Lonstein 2016). These adaptations are underpinned by extensive neurobiological changes that enhance maternal motivation, sensory processing, and cognitive functioning. Among these adaptations, cognitive flexibility - the ability to adapt behavioral responses to changing environmental contingencies - and spatial learning have emerged as critical components of successful caregiving (Uddin 2021). These cognitive processes enable mothers to navigate their environment efficiently, locate resources, and respond effectively to the dynamic needs of their offspring. The study of cognitive flexibility has been relatively limited in mothers, despite the likelihood that these functions are critical for maternal responsiveness and parenting quality in both rodents and humans (Leuner & Gould 2010). Thus, the specific mechanisms and extent of behavioural improvements during the postpartum period remain incompletely determined and understood. The IntelliCage system offers a unique platform to investigate these cognitive changes in rodents under semi-naturalistic and socially enriched conditions (Kiryk et al 2020). This automated system enables the monitoring of multiple animals simultaneously and provides a comprehensive analysis of their behavior across various cognitive tasks (Lipp et al 2023). By allowing precise tracking of individual animals, IntelliCage eliminates the need for human intervention, thereby minimizing stress and enabling the assessment of spontaneous behavior. It also facilitates the implementation of sophisticated behavioral paradigms that probe different dimensions of cognition, including spatial learning, reward-based motivation, and cognitive flexibility (Bramati et al 2023), particularly through paradigms such as place learning and reversal learning (Voikar et al 2018). Place learning tasks assess an animal’s ability to associate a specific location with a reward (Diviney et al 2013), while reversal learning evaluates cognitive flexibility by requiring animals to adapt to a new reward location (Izquierdo et al 2017). These paradigms are particularly relevant for studying maternal cognitive adaptations, as they reflect the types of challenges mothers face in the wild, such as finding food, avoiding predators, and managing the competing demands of caregiving and self-maintenance.

Although cognitive enhancements during the postpartum period have been documented in other experimental setups (Cost et al 2014), no studies have systematically examined these changes in the context of the IntelliCage system. Moreover, existing studies often focused on either the physiological or behavioral aspects of maternal adaptations, with limited integration of these perspectives. Addressing this gap, the present study aims to provide a comprehensive analysis of cognitive flexibility and spatial learning in postpartum females using IntelliCage.

The present study employed a sequence of behavioral tasks designed to assess multiple dimensions of cognition and motivation across reproductive states: pregnancy, postpartum, and control (non-pregnant) conditions. During the adaptation phases, animals were introduced to the IntelliCage environment and trained to perform nosepoke behaviors to access water. Following this, place learning and reversal learning tasks were conducted to evaluate spatial learning and cognitive flexibility. These tasks required animals to associate specific corners with water availability and adapt to changes in reward contingencies. Finally, fixed schedule drinking tasks were implemented to examine anticipatory and persistent behaviors in response to restricted water access.

## Methods

### Animals

The study was approved by the Workplace Animal Welfare Committee of the National Scientific Ethical Committee on Animal Experimentation at Eötvös Loránd University in Budapest, with protocol code PE/EA/568-7/2021. The research involving mice was conducted in accordance with the standards set forth by the Animal Hygiene and Food Control Department of the Ministry of Agriculture in Hungary (40/2013), which are consistent with the EU Directive 2010/63/EU for animal experiments. The animals were housed in accordance with standard laboratory conditions, maintaining a temperature of 23 ± 1 °C and humidity levels between 50 and 60% both when being in the IntelliCage, and before and after that. They experienced a 12-hour light/dark cycle, with the lights turning on at 5:00 AM. They were provided with unlimited access to food (SM R/M-H, 1534-00; Ssniff, Soest, Germany) and drinking water.

Overall, 44 female C57BJ/6J mice (control n=14, pregnant n=23, mother n=7) were used for this study. At 2-3 months of age, prior to IntelliCage testing, a radiofrequency identification transponder chip was subcutaneously implanted into the dorso-cervical region under isoflurane inhalation anesthesia. Mice were allowed 1 week to recover and were then introduced into the IntelliCage. Place learning began on days 4-5 of pregnancy and days 6-12 of the postpartum period. The onset of reversed place learning coincided with days 9-10 of gestation and days 11-16 of lactation. Fixed schedule drinking started on days 14-15 of pregnancy and days 1-2 of the postpartum period.

### IntelliCage setup

The IntelliCage apparatus, with dimensions of 39 cm × 58 cm × 21 cm, comprised of four accessible corners, accessible via an antenna-equipped tunnel. Scanners situated at each corner entrance were programmed to detect the entry of each animal by reading the previously implanted RFID. The sensors located on the nose port and waterspout, respectively, were employed to detect instances of nosepoking and licking. The mice were provided with unlimited standard mouse chow in the middle compartment of IntelliCage throughout the entire experiment. The lights in the behavioral room were kept on from 05:00 to 17:00.

### Data acquisition and visualization

The following data was recorded and obtained from the IntelliCage system: the number of visits, nosepokes and licks made by all animals in all corners. The data obtained from the IntelliCage system was subsequently visualised using bespoke software written in VBA (Visual Basic for Applications) within Microsoft Excel. The software is designed to directly read text files exported from IntelliCage, namely those pertaining to the environment, visits and nosepokes. To ensure consistency, the data is padded with zeros so that records always commence with a complete light or dark period (12 hours). Analytical data is derived from the raw data at varying resolutions (1, 2, 4, 6, or 12 hours) across a comprehensive range of variables, including Correct Visits, Visits without Licks, and so forth. This approach allows for the capture of both the frequency and duration of events. The analytical data can be displayed in two plots, each comprising two graphs. For each plot, the user is able to select the desired time resolution, the point of origin, and the duration of the time period in question. The graphs can display data for individual animals or a chosen average. For averaging, animals can be individually selected (e.g. by sex), and the start time for each animal can be set individually, allowing synchronisation with ovarian cycles, for example. The analytical data displayed on the graphs is also available in table form to facilitate export to external statistical or graphical software. For averages, values for sumX, sumX2, n, and SEM are also included to support further statistical analysis.

### Behavioural tasks

The sequence of behavioral tasks in the IntelliCage was as follows: (1) Adaptation, consisting of free adaptation (1 day) and nosepoke adaptation (3 day) phases; (2) Place learning and reversed place learning; and (3) Fixed schedule drinking (**Figure 1**). The experiments were performed in an IntelliCage apparatus in 4 cycles with all three groups present. Adaptation phase started on days 2-5 of gestation and days 3-6 of lactation. Place learning began on days 4-5 of pregnancy and days 6-12 of the postpartum period. The onset of reversed place learning coincided with days 9-10 of gestation and days 11-16 of lactation. Fixed schedule drinking started on days 14-15 of pregnancy and days 1-2 of the postpartum period.

**Figure 1.**
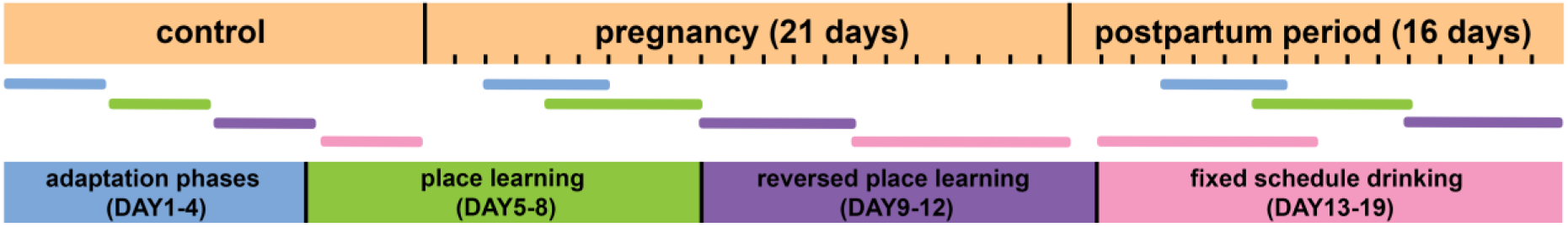
Behavioral battery timeline. The different colored lines indicate the individual tasks and their positioning indicates the start and duration of the task according to the reproduction cycle. Because the length of the overall tasks is collectively longer than the given reproduction cycle, overlaps imply that another group in the same reproduction cycle has performed the task.

Overall data come from a total of 3 experiments. In the first experiment, the number of control animals was 5 and the number of pregnant females was 11. The animals were mated before they entered the IntelliCage and gave birth before the fixed schedule drinking task, so they were already in the early postpartum period in this task. In the second experiment, 12 mice participated (n=5 for control, and 7 for mother group, respectively). The mothers had already given birth before they were transferred to IntelliCage. In the third experiment, as in the first experiment, animals were mated before entering the IntelliCage. In this experiment, 4 control animals and 12 pregnant animals completed the tasks. Data from the experiments were averaged across groups.

#### (1) Adaptation phases

Free adaptation:

During the first day, all doors were left open, allowing the mice unrestricted access to the water bottles, helping them acclimate to the new environment. The following metrics were calculated for this phase:

Total visits: measured to assess exploratory behavior.

Total nosepokes: measured to assess appetitive motivation.

Total licks: measured to assess water consumption.

Visits with lick %: measured to assess water-seeking motivation.

Visits with nosepoke %: measured to assess learning abilities.

Average lick number/ nosepoke: measured to assess consummatory behavior.

Nosepoke adaptation:

In the next four days, the doors remained closed and could be opened only with a nosepoke in a corner, training the animals to nosepoke to access water. In addition to total visits and total licks, the following metrics were also calculated:

Number of Visits with ≥1 Nosepoke Over Total Visits to indicate adaptation and learning. Number of Visits with ≥1 Lick Over Total Visits to indicate the number of visits driven by water-seeking motivation rather than exploratory behavior.

#### (2) Place learning and reversed place learning

Place learning phase:

During this phase, water was accessible in only one of the four corners for each mouse. The specific corner for each mouse was chosen based on their previous visit habits, selecting from the least visited corners to avoid any inherent corner bias. To prevent overcrowding and learning by imitation, the designated reward corners were balanced by the number of mice, with a limit of four mice per corner. For the four days duration of the place learning phase, water was available exclusively in the designated reward corner. Besides measuring total visits and licks per day, the following calculations, called „Correct%” were performed for this task: (Number of visits to the correct corner with a nosepoke) / (Total corner visits). Preference% in place learning phase represents the difference in correct% measured between the final day of nosepoke adaptation and the initial day of place learning. In addition, the slope of the Correct% time curve was measured to asses the speed of learning during the investigated period.

Reversed place learning phase:

For the following four days, water was available only in the opposite corner (reversal). The same parameters were recorded as for the place learning phase.

#### (3) Fixed schedule drinking

For the subsequent seven days, water access was restricted to two 1-hour periods during the light phase (starting at 10 AM and 3 PM). The animals’ access to water was limited only in time, not in space, i.e. they could drink in all four corners. Mice still had to nosepoke to open access to the spout, but any nosepoke outside these periods did not grant access to the water. The behavior of the mice was analyzed 1 hour before, during, and 1 hour after water access to assess anticipatory and persistent behaviors.

The following metrics were calculated in this phase:

Total visits

Total nosepokes

Total licks (only during water access)

### Statistical Analyses

The IntelliCage data were analyzed using 2-way ANOVA with repeated measures when applicable, employing GraphPad Prism 8.1.2. One factor was the group of animals, the other was the time (4 hours or days). When significant changes were found, Bonferroni’s corrected post hoc tests were conducted. A significance level of p<0.05 was used.

## Results

### Adaptation phase

Before facing any challenge, it is crucial to assess the animals’ baseline activity and their responses to the new environment. During the adaptation, the number of total visits decreased across days in all groups (**Figure 2A**) interpreted as decreased exploratory activity. We found a significant reproductive status by day interaction (F(6,69) = 6.067, p<0.0001); post hoc analysis revealed that pregnant mice made more corner visits than control and mother mice on day 1 [control vs. pregnant: ****p<0.0001; control vs mother: ns (non-significant); pregnant vs. mother: ****p<0.0001) and day 2 (control vs. pregnant: *p<0.05, control vs mother: ns, pregnant vs. mother: ***p<0.001]. The number of total nosepokes also decreased over time (F(3,69) = 37.23) (day1: control vs. pregnant: ns; control vs. mother: *p<0.005; pregnant vs. mother: ns) (**Figure 2B**) suggesting decreased appetitive motivation.

**Figure 2.**
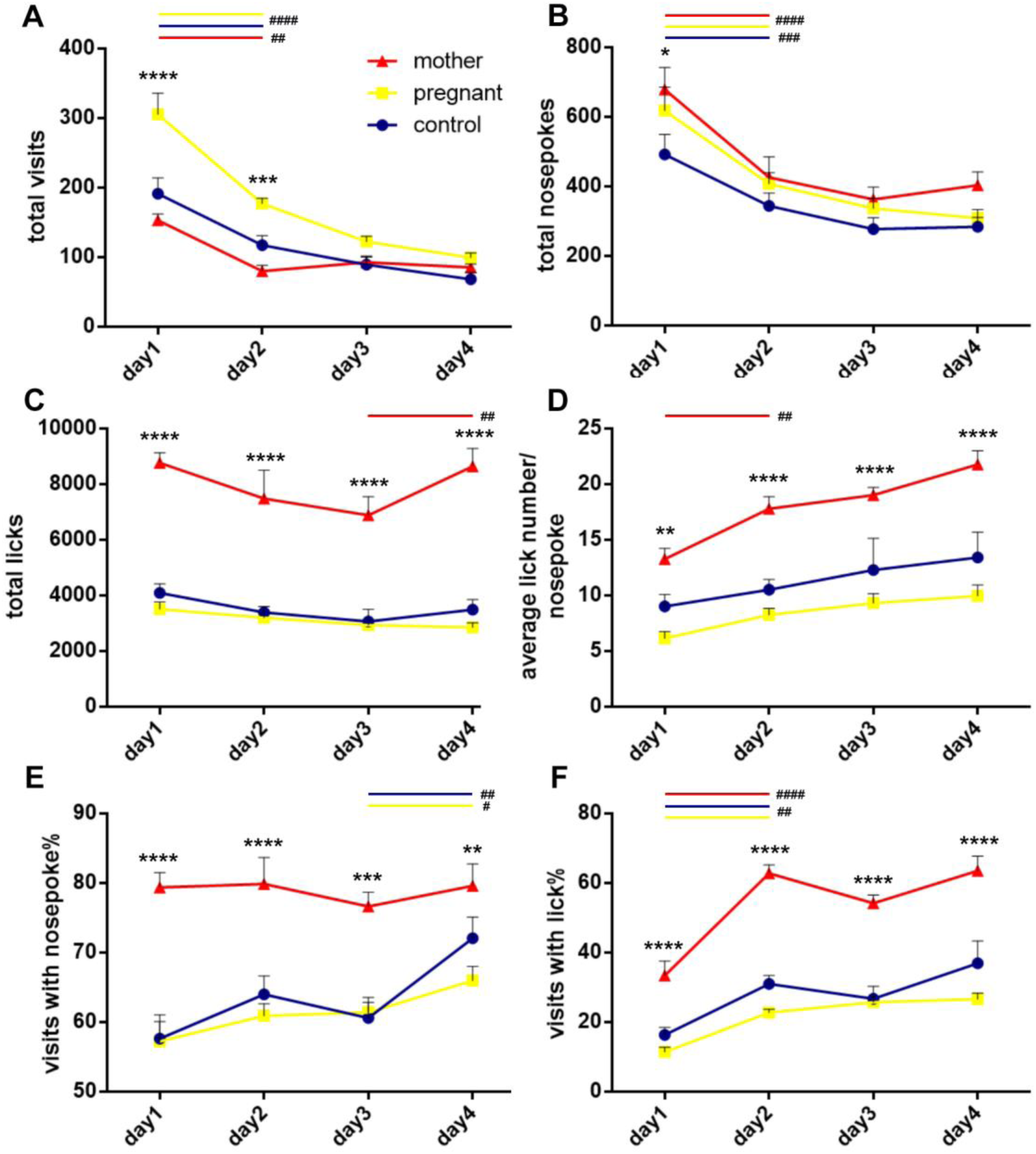
Adaptation phase. Throughout the 4-day adaptation period, mice were able to move freely into any of the four corners and had access to water. (A) We found that pregnant mice (in early pregnancy) made more corner visits on day 1 and 2 compared to mother (in early postpartum period) and control mice. We also found that the number of visits decreased from day 1-4 in all groups. (C) Significant differences were detected in the number of total licks throughout the entire period between mother mice and pregnant/control mice. Nosepoke adaptation phase started on 2nd day, when animals must learn to nosepoke to access water. (E) From day 1, significant differences in visits with nosepoke% were observed between mother and pregnant/control mice. (F) We found that mother mice made more total visits with lick than pregnant and control mice from day 1 of adaptation. Data are presented as means +/- SEM. **p<0.01, ***p<0.001, ****p<0.0001.

Significant differences (control vs. mother and pregnant vs. mother: ****p<0.0001 on every day of the adaptation phases) in total licks were found (F(2,23) = 77.33) (**Figure 2C**), indicating that water consumption varied between mother mice and control/pregnant females. From day 1, significant differences in visits with nosepoke% were observed between mother and control/pregnant mice (F(2,23) = 9.776) (day 1: control vs. pregnant: ns, control vs mother: ****p<0.0001, pregnant vs. mother: ****p<0.0001; day 2: control vs. pregnant: ns, control vs mother: ***p<0.001, pregnant vs. mother: ****p<0.0001; day 3: control vs. pregnant: ns, control vs mother: ***p<0.001, pregnant vs. mother: ***p<0.001; day 4: control vs. pregnant: ns, control vs mother: ns, pregnant vs. mother: **p<0.01) (**Figure 2D**), indicating better learning abilities for mothers. Approximately 40-60% of the mother’s corner visits included drinking behaviour, illustrating increased water-seeking motivation to enter a corner, while visits with lick% was significantly lower in control/pregnant mice (F(2,23) = 67.23) (day 1: control vs. pregnant: ns, control vs mother: ***p<0.001, pregnant vs. mother: ****p<0.0001; day 2: control vs. pregnant: ns, control vs mother: ****p<0.0001, pregnant vs. mother: ****p<0.0001; day 3: control vs. pregnant: ns, control vs mother: ****p<0.0001, pregnant vs. mother: ****p<0.0001; day 4: control vs. pregnant: *p<0.05, control vs mother: ****p<0.0001, pregnant vs. mother: ****p<0.0001) (**Figure 2E**).

Mother mice performed significantly greater numbers of licks per nosepoke (F(2,23) = 18.97) (day 1: control vs. pregnant: ns, control vs mother: *p<0.05, pregnant vs. mother: ns; day 2: control vs. pregnant: ns, control vs mother: ns, pregnant vs. mother: *p<0.05; day 3: control vs. pregnant: ns, control vs mother: *p<0.05, pregnant vs. mother: **p<0.01; day 4: control vs. pregnant: ns, control vs mother: **p<0.01, pregnant vs. mother: ****p<0.0001), indicating increased consummatory behaviour (**Figure 2F**).

A significant decrease in total visits and total nosepokes was observed between day 1 and day 2 in all three groups, which did not show a significant decrease afterwards, indicating that all groups had adapted to the new environment during the 4 days (control, pregnant: day1 vs. day2: ^####^p<0.0001, mother: day1 vs. day2: ^##^p<0.01) (**Figure 2A, B)**. There was also a significant difference in the average number of licks per nosepoke between day 1 and day 2 in mothers (mother: day1 vs. day2: ^##^p<0.01) (**Figure 2D**). The visits with lick% also increased significantly between day 1 and day 2, which indicates that the animals in all three groups were also accustomed to having access to drinking water in the corners (control, pregnant: day1 vs. day2: ^##^p<0.01, mother: day1 vs. day2: ^####^p<0.0001 (**Figure 2F**).

### Place learning

During place learning phase, we found no significant differences in total visits except the third day (F(8,96) = 5.272) (control vs. pregnant: ns, control vs. mother: ns, pregnant vs. mother: *p<0.05) (**Figure 3A**). Similar to the adaptation phase, number of total licks was significantly different between control/pregnant females and mothers during the entire duration of the learning phase (F(2,24) = 84.72) (control vs. mother and pregnant vs. mother: ****p<0.0001 on every day of the place learning phase). We found a significant main effect of day for correct % in the place learning phase (F(3,75) = 8.412, p<0.0001) (**Figure 3C**), illustrating improved learning performance across the 5 days for all groups. We also established a main effect of reproductive status (F(2,25)=14.86, p<0.0001) where mother mice had a significantly higher correct% than control and pregnant female mice from day 2 to day 4 (day 2: control vs. pregnant: ns, control vs. mother : *p<0.05, pregnant vs. mother: **p<0.01; day 3: control vs. pregnant: ns, control vs. mother: ***p<0.001, pregnant vs. mother: ****p<0.0001; day 4: day 3: control vs. pregnant: ns, control vs. mother : ****p<0.0001, pregnant vs. mother: ***p<0.001), but by day 5 the significant difference between pregnant and mother mice diminished. We examined the learning performance of the groups on the first day at a resolution of 4 h and found that significant differences in correct% are already present from the second half of the first day (F(2,25)=6.596) (4th 4h: control vs. pregnant: ns, control vs. mother : *p<0.05, pregnant vs. mother: ns, 5th and 6th 4h: control vs. pregnant: ns, control vs. mother: ns, pregnant vs. mother: *p<0.05) (**Figure 3D**). Preference% was determined as the difference in correct% measured between the final day of nosepoke adaptation (when mice could still drink from all four corners) and the initial day of place learning (when water was available only in the correct corner) and we also found significant differences between groups in this parameter (F(2,25)=10.34) (control vs. pregnant: ns, control vs. mother: ***p<0.001, pregnant vs. mother: **p<0.01) (**Figure 3D**). **Figure 3F** shows the slope of the correct% curve between day 1 and day 2 with the following significance levels (F(2,25)=5.065): control vs. pregnant: ns, control vs. mother: ns, pregnant vs. mother: *p<0.05.

**Figure 3.**
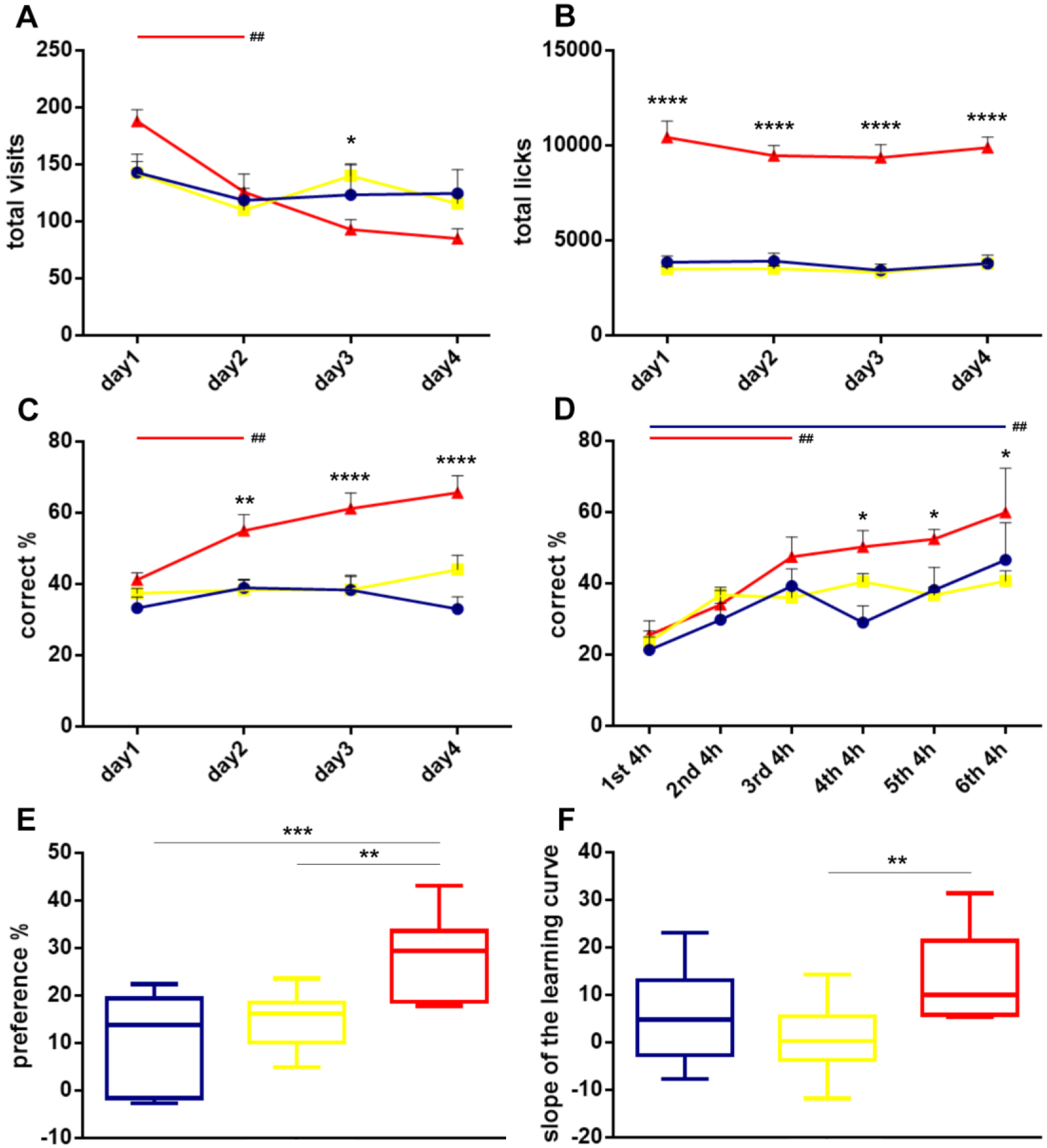
Place learning phase. During the 5-day place learning phase, each animal was assigned to a specific corner where water was available. The other corners were considered incorrect and did not provide access to water. (A) Only on day 3, we found a significant difference in total visits between mother (in mid-postpartum) and pregnant (in early pregnancy) mice. (B) Significant differences in total licks across the 5 days were detected between mother and control/pregnant mice. (C) We found a significant effect of day for correct %, illustrating learning across the 5 days, moreover, significant differences were detected from day 2 between mother and control mice. (D) Significant differences between mothers and control/pregnant mice are already observed in the fourth 4 hours of the day 1 in correct%. (E) Significant differences were detected in preference % between mother and control/pregnant mice. Data are presented as means +/- SEM. (F) The slope of the learning curve of the mothers between day 1 and day 2 also differed significantly from that of pregnants. *p<0.05, **p<0.01, ***p<0.001, ****p<0.0001.

During this phase, total visits reduced, but correct% increased significantly between day 1 and day 2 in mothers (mother: day1 vs. day2: ^##^p<0.01) (**Figure 3A, C**). The percentage of correct corner visits (correct%) measured on the first day was already significantly increased between 1st and 3rd 4h in mothers, while it was significantly increased between 1st and 6th 4h in control group (F(2,25)=6.596) (control: 1st 4h vs. 6th 4h: ^##^p<0.01, mother: 1st 4h vs. 3rd 4h: ^##^p<0.01) (**Figure 3D**).

### Reversed place learning

Next, animals were assessed in the reversed place learning phase, where the correct corner was opposite to that of the first phase of place learning. We found significant differences in total visits only on day 1 (control vs. pregnant: ns, control vs. mother: ns; pregnant vs. mother: **p<0.01) and day 2 (control vs. pregnant: ns, control vs. mother: *p<0.05; pregnant vs. mother: ****p<0.0001) (F(2,24)=4.815) (**Figure 4A**). Water consumption between control/mother mice and pregnant female mice was not significantly different (**Figure 4B**). For correct%, we found a significant main effect of day (F(3,75)=10.64, p<0.0001; **Figure 4C**), illustrating learning across the 5 days. Additionally, we also found that the mother mice had a significantly higher correct% from the second day compared to control and pregnant mice (F(2,25) = 14.70, p<0.0001; day 2: control vs. pregnant: ns, control vs. mother : **p<0.01, pregnant vs. mother: ****p<0.0001; day 3: control vs. pregnant: ns, control vs. mother : **p<0.01, pregnant vs. mother: ****p<0.0001; day 4: control vs. pregnant: ns, control vs. mother : ****p<0.0001, pregnant vs. mother: ****p<0.0001; day 5: control vs. pregnant: ns, control vs. mother : ****p<0.0001, pregnant vs. mother: ***p<0.001) (**Figure 4C**). We also examined the learning performance of the animals on the first day of this test at a resolution of 4 hours, and found that the correct% was again significantly different for the mothers compared to the control and pregnant females from the 3rd 4h (F(2,25)=9.826) (3rd 4h: control vs. pregnant: ns, control vs. mother : ns, pregnant vs. mother: *p<0.05; 4th 4h: control vs. pregnant: ns, control vs. mother : **p<0.01, pregnant vs. mother: *p<0.05; 5th 4h: control vs. pregnant: ns, control vs. mother: ns, pregnant vs. mother: ***p<0.001; 6th 4h: control vs. pregnant: ns, control vs. mother:*p<0.05; pregnant vs. mother: ***p<0.001) (**Figure 4D**). In this case, preference% represents the difference of correct% between the last day of place learning and the first day of reversed place learning and there is a significant difference between control mice and mother mice (F(2,25)=6.142) (control vs. mother: **p<0.01) (**Figure 4E**). Figure 4F shows the slope of the learning curve between day 1 and day 2, where a significant difference was observed only between the slope of the learning curve of mothers and pregnants (F(2,25)=6.089) (pregnant vs. mother: **p<0.01).

**Figure 4.**
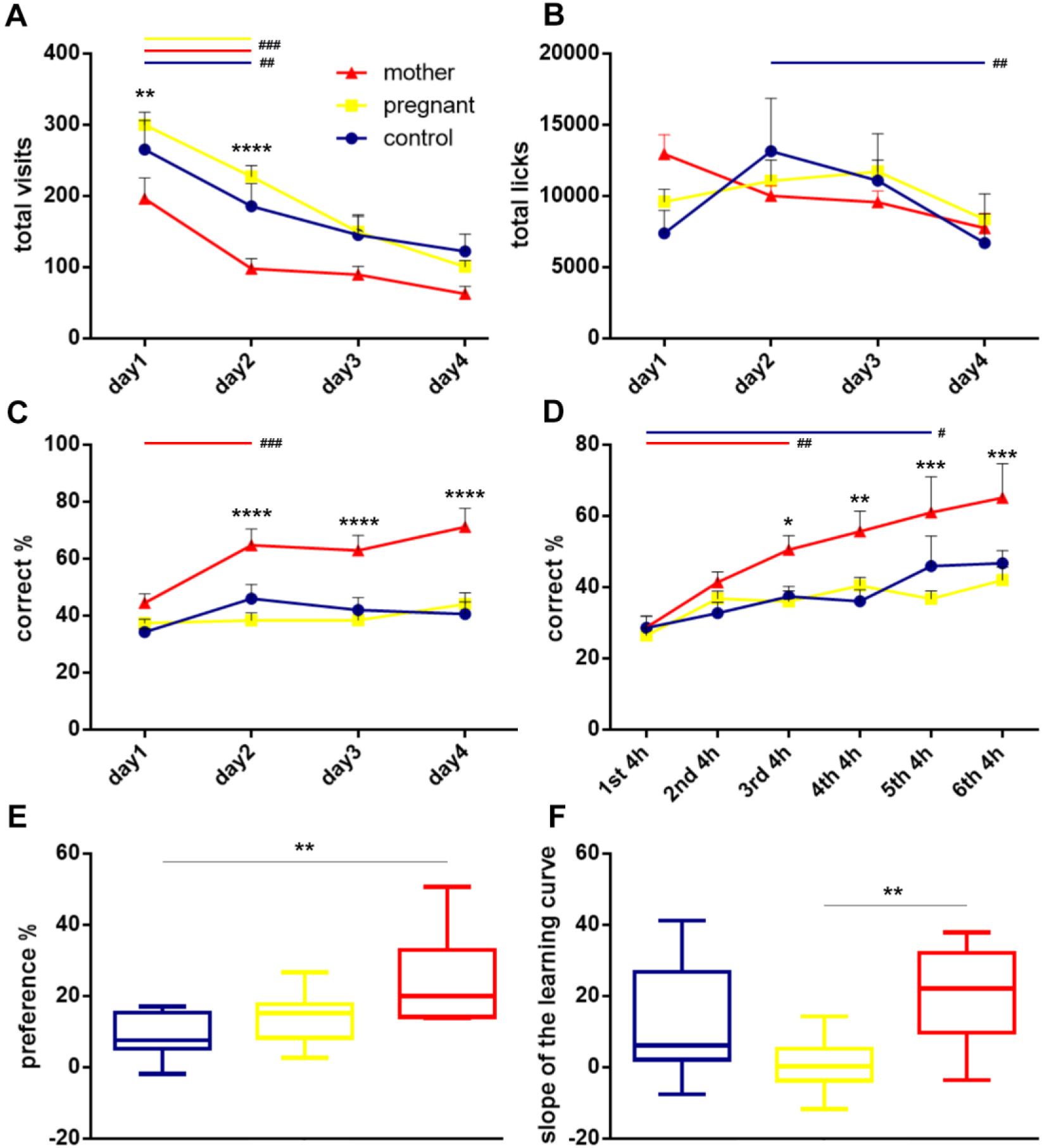
Reversed place learning phase. During the reversed place learning phase, animals could obtain water by entering and nose-poking the corner opposite to the one assigned during the initial place learning phase. (A) Significant differences in total visits were detected on day 1 and day 2 between pregnant (in mid-pregnancy) and mother (in late postpartum) mice. (B) No differences in the total number of licks over the 5 days were observed. (C) We found that mother mice made a higher percentage of visits to the correct corner from day 2 than control/pregant mice. Additionally, correct% increased across the 5 days in all groups, illustrating learning. (D) We found that mother mice had a higher correct% than control/pregnant mice from the third 4 hours of the first day. (E) There were also significant differences in preference % between control females and mothers. (F) The slope of the learning curve of the mothers between day 1 and day 2 also differed significantly from that of pregnants. Data are presented as means +/-SEM. *p<0.05, **p<0.01, ***p<0.001, ****p<0.0001.

### Fixed schedule drinking

We focused on the one-hour periods before, during and after the access to water in order to analyze anticipatory and persistent behavior of mice. The number of visits increased in mother mice during the hours preceding water access as the days progressed, while for the other two groups, there was no similar change (1. drinking session: (F(12,174)=5.345), day 2: control vs. pregnant: ns, control vs. mother: ns, pregnant vs. mother: *p<0.05; day 4: control vs. pregnant: ns, control vs. mother: *p<0.05, pregnant vs. mother: **p<0.01; day 6: control vs. pregnant: ns, control vs. mother : ***p<0.001, pregnant vs. mother: *p<0.05; day 7: control vs. pregnant: ns, control vs. mother: ****p<0.0001, pregnant vs. mother: ****p<0.0001 (**Figure 5A**), 2. drinking session: (F(12,240)=2.335), day 4: control vs. pregnant: *p<0.05, control vs. mother: ns, pregnant vs. mother: ns; day 6: control vs. pregnant: ns, control vs. mother : *p<0.05, pregnant vs. mother: ns; day 7: control vs. pregnant: ns, control vs. mother: ****p<0.0001, pregnant vs. mother: ****p<0.0001) (**Figure 5B**). During the hours when water was accessible, the number of visits showed significant differences between mother mice and control/pregnant mice (1. drinking session: (F(12,204)=3.435), day 1: control vs. pregnant: **p<0.01, control vs. mother: ns, pregnant vs. mother: *p<0.05; day 5: control vs. pregnant: ns, control vs. mother: **p<0.01, pregnant vs. mother: ***p<0.001; day 6: control vs. pregnant: ns, control vs. mother: ns, pregnant vs. mother: ***p<0.001; day 7: control vs. pregnant: ns, control vs. mother: ****p<0.0001, pregnant vs. mother: ****p<0.0001 (**Figure 5C**), 2. drinking session: (F(12,240)=3.366), day 2: control vs. pregnant: ns, control vs. mother: **p<0.01, pregnant vs. mother: **p<0.01; day 6: control vs. pregnant: ns, control vs. mother : **p<0.01, pregnant vs. mother: **p<0.01; day 7: control vs. pregnant: ns, control vs. mother : **p<0.01, pregnant vs. mother: **p<0.01) (**Figure 5D**). During the hours following water access, lower visits for mothers were observed as the days progressed compared to control/pregnant mice (1. drinking session: (F(12,174)=1.276), day 1: control vs. pregnant: ns, control vs. mother: *p<0.05, pregnant vs. mother: ns (**Figure 5E**), 2. drinking session: (F(12,240)=2.854), day 1: control vs. pregnant: ns, control vs. mother: ***p<0.001, pregnant vs. mother: ns; day 2: control vs. pregnant: ns, control vs. mother : **p<0.01, pregnant vs. mother: **p<0.01) (**Figure 5F**). So after four days, mother mice increased their number of visits during the hours preceding water access, which may be interpreted as an anticipation of the up-coming event (water access) and did not show persistent higher activity after water was not available anymore.

**Figures 5.**
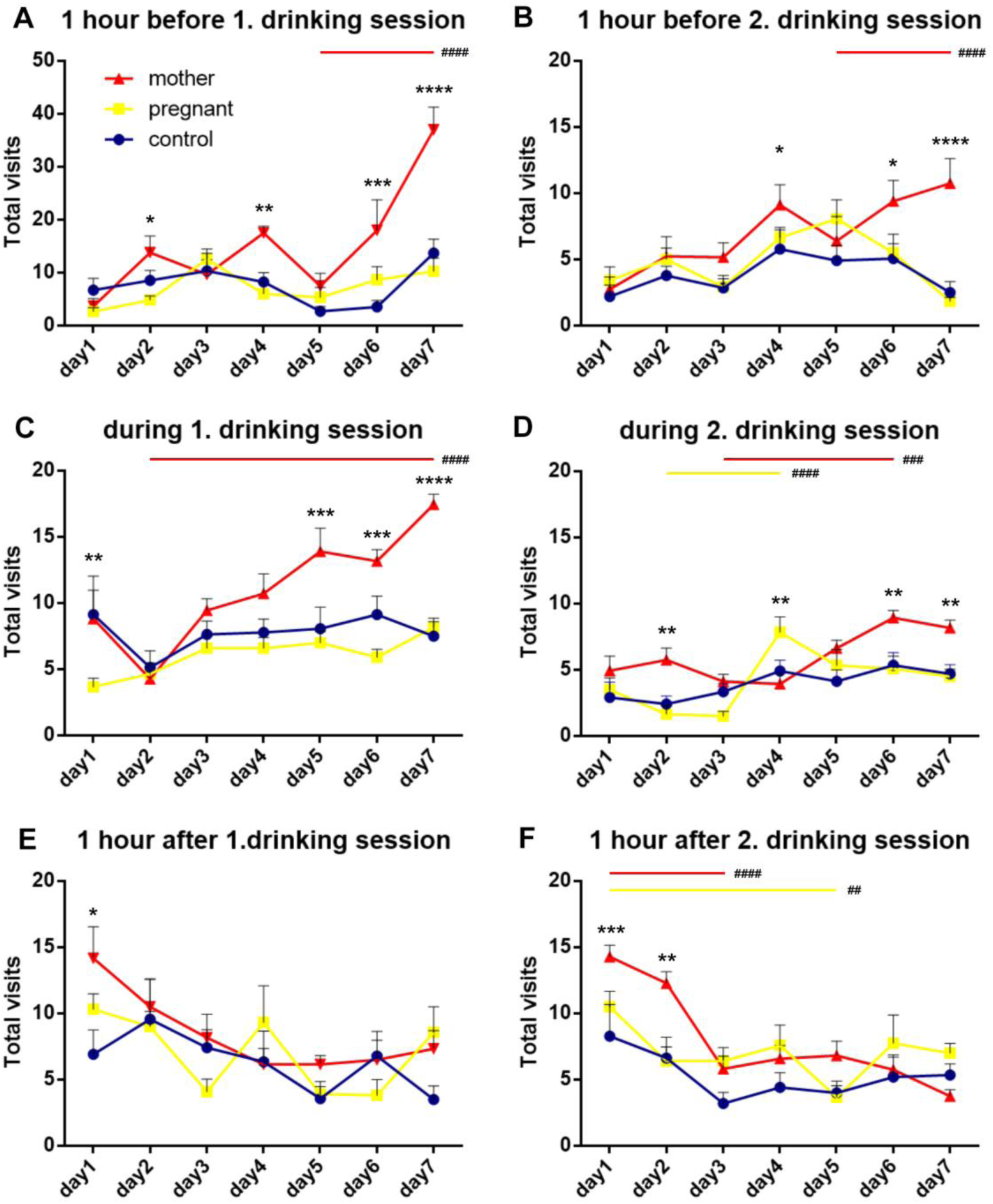
Total visits in fixed schedule drinking phase. Number of visits 1 hour before water accessibility (A, B), during water access (C, D) and 1 hour after water accessibility (E, F). Pregnant mice were in late pregnancy and mother mice were in early postpartum period for this task. Data are presented as means +/- SEM. *p<0.05, **p<0.01, ***p<0.001, ****p<0.0001.

Significant differences were found between day 5 and day 7 in the hours before drinking sessions only for mothers (mother: day5 vs. day7 before 1. and 2. drinking session: ^####^p<0.0001) (**Figure 5 A, B)**. The number of visits during drinking sessions also showed a significant increase in mothers between day 2 and day 7 (during 1. drinking session) and between day 3 and day 6 (during 2. drinking session) (mother: day2 vs. day7 during 1. drinking session: ^####^p<0.0001; mother: day3 vs. day6 during 2. drinking session: ^###^p<0.001) (**Figure 5 C, D)**. In the hour following the 2. drinking session, a significant effect of the days was found between day 1 and day 3 in the mothers and between day 1 and day 5 in the pregnant group (mother: day1 vs. day3: ^####^p<0.0001; pregnant: day1 vs. day7: ^##^p<0.01) (**Figure 5F**). It is also important to note that we did not find any increase in the mortality of pups during the procedure.

The number of nosepokes exhibited a proportional relationship with the total number of visits, resulting in curves that were similar to those observed for total visits with the following significance levels: 1 hour before 1. drinking session: F(12,174)=3.084, day 6: control vs. pregnant: ns, control vs. mother : ****p<0.0001, pregnant vs. mother: ***p<0.001; day 7: control vs. pregnant: ns, control vs. mother: ***p<0.001, pregnant vs. mother: **p<0.01 (**Figure 6A**), 1 hour before 2. drinking session: F(12,240)=0.7662, day 4: control vs. pregnant: ns, control vs. mother: *p<0.05, pregnant vs. mother: *p<0.05; day 7: control vs. pregnant: ns, control vs. mother: *p<0.05, pregnant vs. mother: ns (**Figure 6B**), during 1. drinking session: F(12,204)=2.910, day 1: control vs. pregnant: *p<0.05, control vs. mother : ns, pregnant vs. mother: ns; day 3: control vs. pregnant: **p<0.01, control vs. mother: ns, pregnant vs. mother: ns; day 5: control vs. pregnant: ns, control vs. mother: *p<0.05, pregnant vs. mother: **p<0.01; day 6: control vs. pregnant: ns, control vs. mother: ns, pregnant vs. mother: ****p<0.0001; day 7: control vs. pregnant: ns, control vs. mother: *p<0.05, pregnant vs. mother: *p<0.05 (**Figure 6C**), during 2. drinking session: F(12,204)=4.641, day 6: control vs. pregnant: ns, control vs. mother: ****p<0.0001, pregnant vs. mother: ****p<0.0001; day 7: control vs. pregnant: ns, control vs. mother: ***p<0.001, pregnant vs. mother: ***p<0.001 (**Figure 6D**), 1 hour after 1. drinking session: F(12,174)=1.981, day 2: control vs. pregnant: *p<0.05, control vs. mother: ns, pregnant vs. mother: **p<0.01; day 3: control vs. pregnant: ns, control vs. mother: ns, pregnant vs. mother: **p<0.01 (**Figure 6E**), 1 hour after 2. drinking session: F(12,174)=2.603, day 1: control vs. pregnant: ns, control vs. mother: **p<0.01, pregnant vs. mother: ns; day 2: control vs. pregnant: ***p<0.001, control vs. mother: ns, pregnant vs. mother: ns (**Figure 6F**). So not only the exploratory activity of the mothers increased during the hours preceding water access, but also the number of nosepokes specifically associated with drinking.

**Figures 6.**
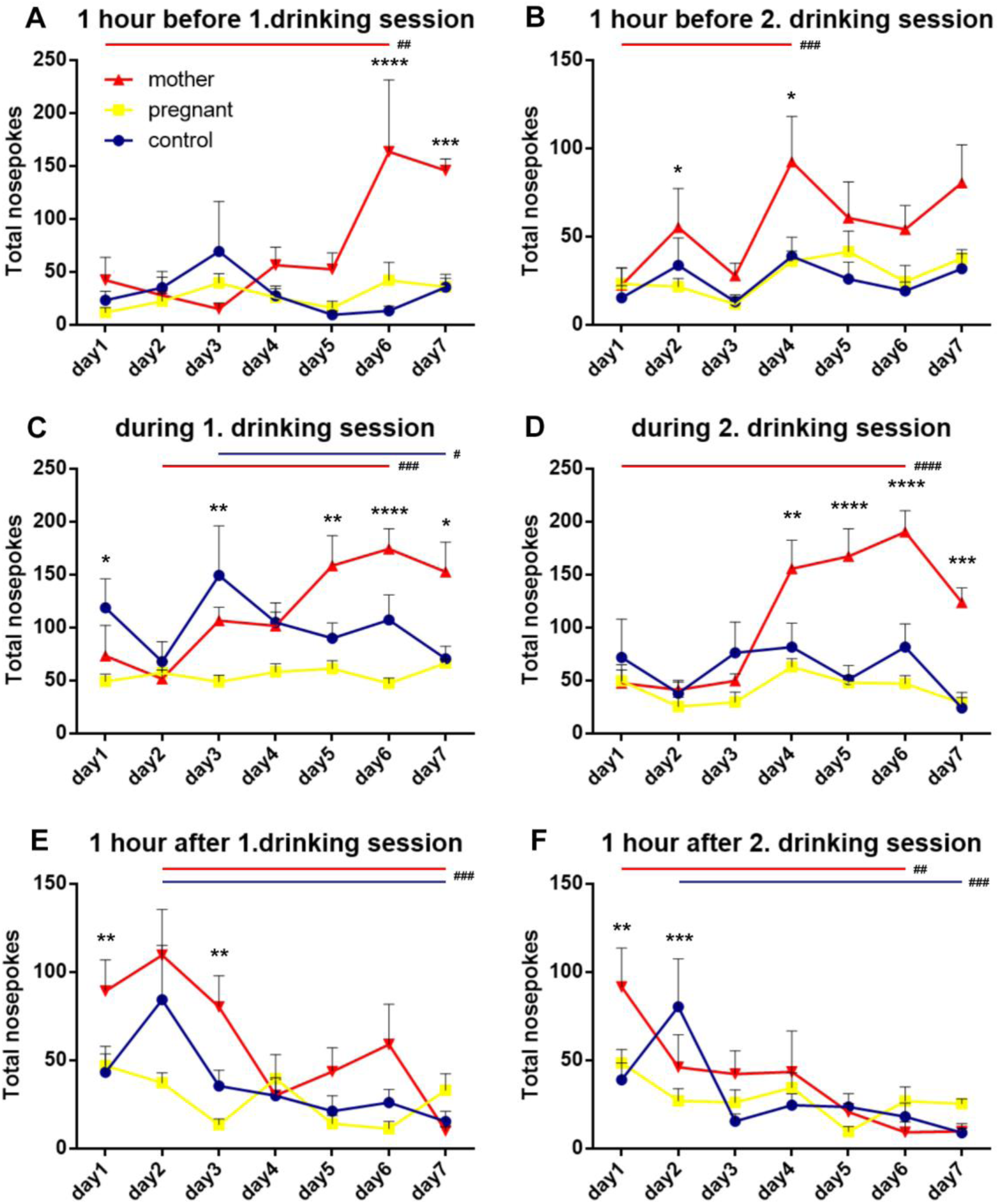
Total nosepokes in fixed schedule drinking phase. Number of visits 1 hour before water accessibility (A, B), during water access (C, D) and 1 hour after water accessibility (E, F). Pregnant mice were in late pregnancy and mother mice were in early postpartum period for this task. *p<0.05, **p<0.01, ***p<0.001, ****p<0.0001.

In the hours before drinking sessions, the number of nosepokes increased significantly in mothers as the days progressed (mother: day1 vs. day6 before 1. drinking session: ^##^p<0.01; mother: day1 vs. day4 before 2. drinking session: ^###^p<0.001) (**Figure 6A, B)**. We also observed similar effect during drinking sessions (mother: day2 vs. day6 during 1. drinking session: ^###^p<0.001; pregnant: day3 vs. day7 during 1. drinking session: ^#^p<0.1; mother: day1 vs. day6 during 2. drinking session: ^####^p<0.0001) (**Figure 6C, D)**. In the hours following drinking sessions, the number of nosepokes significantly decreased after days in both mothers and controls (mother, pregnant: day2 vs. day7 after 1. drinking session: ^###^p<0.001; mother: day1 vs. day6 during 2. drinking session: ^##^p<0.01; control: day2 vs. day7 during 2. drinking session: ^###^p<0.001) (**Figure 6E, F)**.

During drinking session, there were significant differences in total licks between mothers and control/pregnant mice with the following significance levels: during 1. drinking session: F(12,240)=1.195, day 4: control vs. pregnant: ns, control vs. mother: ns, pregnant vs. mother: *p<0.05; day 5: control vs. pregnant: ns, control vs. mother: **p<0.01, pregnant vs. mother: **p<0.01; day 6: control vs. pregnant: ns, control vs. mother: **p<0.01, pregnant vs. mother: ****p<0.0001; day 7: control vs. pregnant: ns, control vs. mother: **p<0.01, pregnant vs. mother: ns (**Figure 7A**), during 2. drinking session: F(12,240)=1.633, day 2: control vs. pregnant: ns, control vs. mother: **p<0.01, pregnant vs. mother: **p<0.01; day 5: control vs. pregnant: ns, control vs. mother: **p<0.01, pregnant vs. mother: *p<0.05; day 6: control vs. pregnant: ns, control vs. mother: ns, pregnant vs. mother: *p<0.05 (**Figure 7B**). The average lick number per nosepoke also differs significantly between groups, but in this case in favour of pregnant mice [during 1. drinking session: F(12,240)=4.344, day 1: control vs. pregnant: **p<0.01, control vs. mother: ns, pregnant vs. mother: **p<0.01; day 3: control vs. pregnant: **p<0.01, control vs. mother: ns, pregnant vs. mother: **p<0.01; day 5: control vs. pregnant: *p<0.05, control vs. mother: ns, pregnant vs. mother: ns; day 6: control vs. pregnant: *p<0.05, control vs. mother: ns, pregnant vs. mother: ns (**Figure 7C**), during 2. drinking session: F(12,240)=3.245, day 2: control vs. pregnant: *p<0.05, control vs. mother: ns, pregnant vs. mother: *p<0.05; day 5: control vs. pregnant: *p<0.05, control vs. mother: ns, pregnant vs. mother: *p<0.05; day 6: control vs. pregnant: *p<0.05, control vs. mother: ns, pregnant vs. mother: *p<0.05 (**Figure 7D**)].

**Figure 7.**
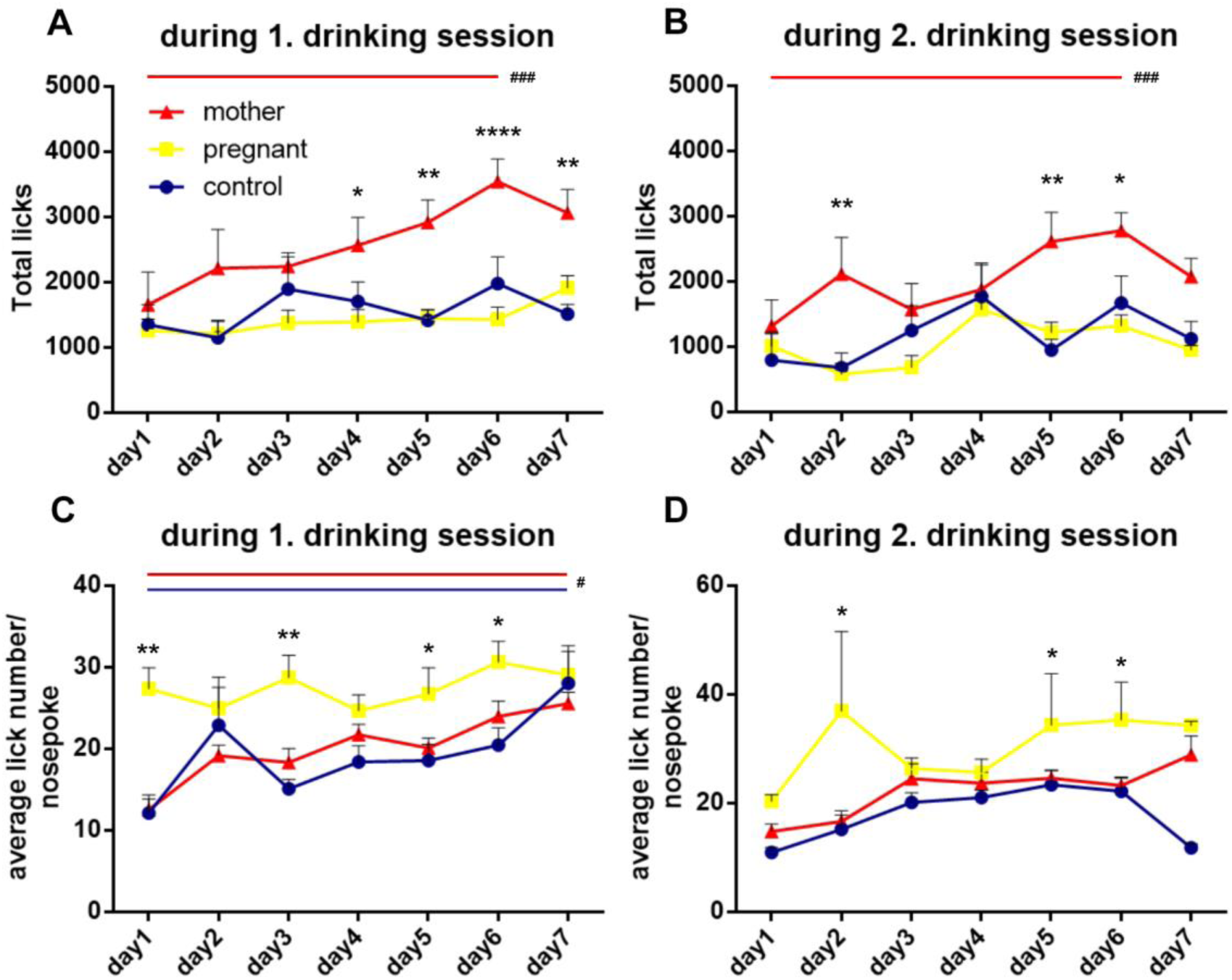
Drinking sessions in fixed schedule drinking phase. Number of licks during the first one-hour (from 10 AM) (A) and second one-hour (from 3 PM) (B) drinking sessions. Average lick number per nosepoke during the first one-hour (from 10 AM) (C) and second one-hour (from 3 PM) (D) drinking sessions. *p<0.05, **p<0.01, ***p<0.001, ****p<0.0001.

In drinking sessions, not only the number of nosepokes but also the number of licks showed a significant increase with the progression of days in mothers (mother: day1 vs. day 6 during 1. and 2. drinking session: ^###^p<0.001 (**Figure 7A, B)**. Days had a significant effect on the number of licks per nosepoke in the first drinking session in mothers and control mice (mother, control: day1 vs. day7 during 1. drinking session: ^#^p<0.1) (**Figure 7C, D)**.

Mother mice modified their corner visit activity after 4 days, as the difference between dark and light periods reversed (**Figure 8A, B)**, with significant changes in the number of nose-pokes as well (**Figure 8C, D)**. Mother mice decreased their number of corner visits during the dark phase (F(12,192)=5.948, day 1: control vs. pregnant: **p<0.01, control vs. mother: ns, pregnant vs. mother: *p<0.05; day 2: control vs. pregnant: ns, control vs. mother: **p<0.01, pregnant vs. mother: ****p<0.0001; day 3: control vs. pregnant: *p<0.05, control vs. mother: **p<0.01, pregnant vs. mother: ****p<0.0001; day 4: control vs. pregnant: ns, control vs. mother: ns, pregnant vs. mother: *p<0.05) and increased in the light phase (F(12,204)=7.938, day 1: control vs. pregnant: ns, control vs. mother: ns, pregnant vs. mother: *p<0.05; day 5: control vs. pregnant: ns, control vs. mother: *p<0.05, pregnant vs. mother: ns; day 6: control vs. pregnant: ns, control vs. mother: ****p<0.0001, pregnant vs. mother: ****p<0.0001; day 7: control vs. pregnant: ns, control vs. mother: ****p<0.0001, pregnant vs. mother: ***p<0.001). However, a marked difference remained between dark and light periods. The same happened with nosepokes around day 4, with the previous higher number of nosepokes during dark phase (F(12,192)=3.173, day 1: control vs. pregnant: ns, control vs. mother: ns, pregnant vs. mother: ***p<0.001; day 2: control vs. pregnant: ns, control vs. mother: ***p<0.001, pregnant vs. mother: ****p<0.0001; day 3: control vs. pregnant: *p<0.05, control vs. mother: ***p<0.001, pregnant vs. mother: ****p<0.0001; day 7: control vs. pregnant: ns, control vs. mother: *p<0.05, pregnant vs. mother: ns) being replaced by a higher number of nosepokes during light phase (F(12,192)=4.244, day 4: control vs. pregnant: ns, control vs. mother: ***p<0.001, pregnant vs. mother: ***p<0.001; day 5: control vs. pregnant: ns, control vs. mother: ***p<0.001, pregnant vs. mother: ***p<0.001; day 6: control vs. pregnant: ns, control vs. mother: ***p<0.001, pregnant vs. mother: ***p<0.001; day 7: control vs. pregnant: ns, control vs. mother: **p<0.01, pregnant vs. mother: **p<0.01). Meanwhile, the number of visits and nosepokes of control/ pregnant females did not change significantly during the 7 days, so they were not able to change their behaviour according to the changed conditions. Overall, control/pregnant mice displayed a significantly lower number of visits and nosepokes during the light periods from day 4 and no increase in these parameters was observed later.

**Figure 8.**
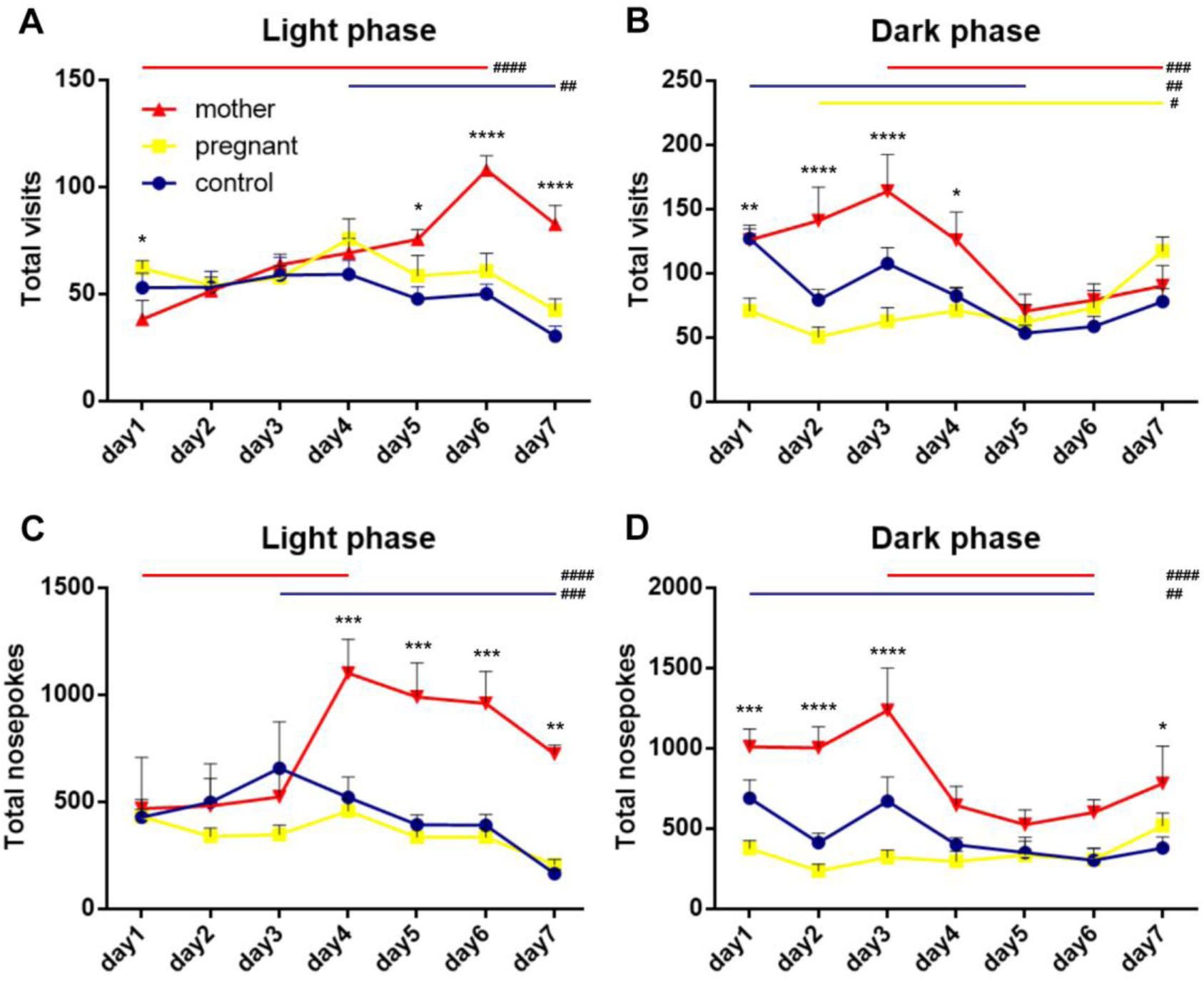
Number of visits and nosepokes in 12-hour bins during fixed schedule drinking. Total number of visits during light phase (A). Total number of visits during dark phase (B). Total number of nosepokes during light phase (C). Total number of nosepokes during dark phase (D).

The number of visits in light phase showed a significant increase in mothers and a significant decrease in controls over the days, while in the dark phase there was a decrease in the number of visits in mothers and control animals and a significant increase in pregnants (mother: day1 vs. day6: ^####^p<0.0001; control: day4 vs. day7: ^##^p<0.01 during light phase; mother: day3 vs. day7: ^###^p<0.001); control: day1 vs. day5: ^##^p<0.01, pregnant: day2 vs. day7: ^#^p<0.1 during dark phase (**Figure 8A, B)**. A similar pattern was observed in the number of nosepokes with the progression of days (mother: day1 vs. day4: ^####^p<0.0001; control: day4 vs. day7: ^##^p<0.01 during light phase; mother: day3 vs. day6: ^####^p<0.0001); control: day1 vs. day6: ^##^p<0.01 during dark phase (**Figure 8C, D)**.

## Discussion

The present study is the first to determine that motherhood improves spatial learning performance and cognitive flexibility using IntelliCage. Previously, standard behavioral tests were employed to examine these phenomena in relation to different reproductive stages, but the results were inconclusive (Albin-Brooks et al 2017)(Tomizawa et al 2003)(Gatewood et al 2005). In turn, the present results revealed significant enhancements in spatial learning and cognitive flexibility in postpartum female mice compared to pregnant and control animals.

### The effect of the reproductive status on different types of learning

During the four-day adaptation period, all animals exhibited signs of acclimatisation to the IntelliCage, as evidenced by frequent visits to all corners and the acquisition of corner-drinking skills through nosepoking. During this process, the initially high visit and nosepoke numbers declined from day 1 to day 2 in all groups and remained stable thereafter. This decline can be interpreted as a result of the elevated exploration observed on the first day of habitation in the IntelliCage, during which the animals familiarised themselves with the corners and nosepoking devices. Concurrently, the total lick numbers exhibited relative stability, attributable to the increase in visits with lick/total visit and the lick number/nosepoke ratios on the second day. This suggests that by that day, the animals had acquired the ability to utilise the nosepoke for obtaining a drink to lick. While these tendencies were evident across all groups, notable variations in behaviour were observed among different groups. A notable observation was the increased exploratory drive exhibited by pregnant mice, as evidenced by their frequent visits to corners, in contrast to the behaviour patterns of the other groups. While no other study has been found that investigates the altered exploratory drive of pregnant animals, the finding of increased exploration could be considered as a preparation for finding a place to build a nest. However, it is important to note that anxiety can also reduce exploratory behaviour. Therefore, reduced anxiety in pregnant animals could also explain the initially increased exploratory behavior in the pregnant group. A further significant disparity between the groups was observed in the elevated licking behaviour exhibited by the postpartum mothers. Given the constant size of the liquid drop in the IntelliCage, the elevated licking number is indicative of increased fluid intake, a phenomenon that is well-supported by the literature and is entirely consistent with the needs of a lactating animal that is responsible for nourishing her offspring. It is important to note that while food is freely available for the animals in the IntelliCage, the amount of water and food intake is proportional due to the strict osmotic regulatory processes if the salt content of the food is constant (Johnson et al 2003), as was the case in our experiment. The increased lick number in mothers was primarily due to an elevated lick number per nosepoke within the 5 seconds during which licking was possible following a nosepoke. However, the number of nosepokes was also somewhat higher in mothers than in the other groups. This finding is significant because it suggests that the number of nosepokes is proportional to the motivation of the appetitive phase of the feeding behaviour (food seeking). In contrast, the intensity of licking is associated with the consummatory, hedonic motivation. These two phases of feeding behaviours are potentially governed by distinct brain mechanisms (Jennings et al 2015). On the first day of the experiment, the animals were suddenly unable to access water in three of the four corners. This led to an increase in the number of visits made by the animals, particularly by the mothers, which decreased by the third day, with the mothers visiting the corners less frequently than the other groups. This decline in exploratory behaviour among mothers is consistent with previous findings in rats (Habr et al 2011).

The elevated lick number observed despite a concomitant reduction in total visitation by mothers can be attributed to the animals’ acquisition of the correct corner for drinking. The ratio of visits to this corner increased twofold in control and pregnant animals, and threefold in mothers during the first 1.5 days. The initial ratio of visits to the correct corners was slightly less than the 25% by chance ratio because the correct corner for each animal was selected to be an originally less preferred one to avoid preference bias. While all groups demonstrated learning of the correct corner, a pronounced difference emerged among the groups. Mothers exhibited a significantly faster rate of improvement in the correct corner ratio compared to the other groups. This finding is further supported by the observation that the difference in correct% measured between the final day of nosepoke adaptation and the initial day of place learning (termed Preference%) was significantly higher in the mother group than in the other groups. Additionally, the mother group exhibited a steeper correct% curve on the first day of place learning. Furthermore, the ratio of visits to the correct corner continued to improve beyond the first day. Consequently, during the process of place learning, postpartum females exhibited a more rapid acquisition of the correct location associated with water and a more pronounced improvement in performance over time. These results are consistent with previous data obtained in the rat, which demonstrated that maternal rats outperform virgin females in spatial tasks. Maternal rats demonstrate a reduced number of errors and exhibit enhanced efficiency in navigating the Morris water maze and object placement tasks. It is noteworthy that these cognitive enhancements are specific to spatial learning, as maternal and virgin rats demonstrate equivalent performance in object recognition (Lemaire et al 2006). In a similar vein, during the phase of reversal learning, the temporal progression of the decline in total visits and the rise in correct corner ratio, along with other attributes of the adaptation learning, exhibited parallels with those observed in place learning. It was also observed that postpartum females demonstrated greater adaptability to the new reward location. The enhanced outcomes in reverse place learning in IntelliCage are generally interpreted as indicative of augmented cognitive flexibility, given its association with the capacity to adapt behaviour in response to altered rules or contingencies, employ alternative strategies when prior ones are no longer effective, and suppress established responses in favour of new ones (Daguano Gastaldi et al 2025)(Ma et al 2023). Therefore, it can be concluded from the findings that mother mice possess superior cognitive flexibility in comparison to both controls and pregnant animals. It is also noteworthy that enhanced spatial learning and cognitive flexibility may confer significant survival advantages by enabling mothers to navigate their environment efficiently, secure resources, and manage the challenges of caregiving.

In the fixed schedule drinking paradigm, mice were provided with water at two-hour intervals, thereby reducing the degree of water restriction and enabling the collection of data twice daily. The distribution of these two hours was not uniform throughout the day; rather, they were separated by a period of five hours (or 19 hours). Consequently, both drinking sessions occurred during the light phase, thereby prompting varying degrees of motivation to drink, owing to the heightened water deprivation following the extended 19-hour interval between sessions. However, the total licks and the number of licks following a nosepoke were not found to be different between the two drinking sessions. However, a subsequent analysis of day-to-day alterations revealed a significant increase in the total licks of the mothers as the schedule progressed, while the other two groups only exhibited a tendency to do so. Of particular interest was the finding that this increase in the total licks of mothers was largely due to an elevation of nosepoking activity between days 3-5, while only a small gradual increase in the number of licks per nosepoke was found in mothers. In contrast, the latter parameter exhibited elevated levels in the pregnant groups from the outset, with the values of the mothers and controls only attaining parity by the seventh day of the schedule. This suggests that the elevated levels of licking per nosepoke observed in mothers (see Adaptation period data) remained constant during the fixed schedule of drinking in mothers, yet it increased similarly or even more so in pregnant and control animals. This finding provides further evidence to support the hypothesis that elevated nose poke is indicative of appetitive drive, and the number of licks per nosepoke is a measure of consummatory drive. The results of this study demonstrate that the increased drinking behaviour of mothers was associated with elevated consummatory drive, suggesting that drinking was more rewarding for them. In contrast, the elevated nose poke and drinking behaviour observed in pregnant animals was not associated with an increase in consummatory drive.

In addition to the drinking sessions, the number of visits and nosepokes was analysed during the hour when the animals had access to water, as well as during the hour before and after this, in order to ascertain whether mice are capable of learning time-based contingencies. It was observed that only the mother mice exhibited an increase in visiting and nosepoking activities prior to the drinking sessions. The elevated exploration levels were observed to commence on day 4 and to increase until day 6. This anticipatory parameter is used to assess temporal learning, and it can be concluded that mothers have a superior cognitive capacity for time tracking in comparison to controls and pregnant subjects. Alternatively, the elevated anticipatory behaviour may be attributable to increased motivation, given that both changes in capability and motivation have the capacity to affect behaviour (Dobolyi 2025).

A marked difference in behaviour was also observed between the groups following the drinking session. Mother mice, and to a lesser extent the control group, persisted in their visits and nosepoking, while the pregnant subjects did not. This behaviour was no longer observed by the third day of the fixed drinking schedule, indicating an effective extinction learning process. An examination of visit and nosepoke activities in periods outside of drinking sessions revealed a significant group difference, too. In the dark phase (when no drinking session was scheduled), mothers exhibited a significantly higher number of visits and nosepokes compared to the control and pregnant animals. A marginal increase in these behaviours was observed in the control group when compared to the pregnant group. These behaviours diminished between days 3 and 5, suggesting extinction learning. Conversely, a marked increase in visit and nosepoke behaviour was observed in mothers during the dark phase (D3-D5), indicating a potential learning process related to the established schedule. In contrast, a decline in visits and nosepokes was observed in the light phase, a phenomenon that was particularly evident in the pregnant and control groups. Thus, fixed schedule drinking tasks highlighted increased anticipatory and persistent behaviors in postpartum females, reflecting heightened motivation. These findings suggest that the postpartum period is associated with enhancements in multiple cognitive domains.

By leveraging the IntelliCage system to study these adaptations, this research contributes to a growing body of literature on the interplay between reproduction and cognition. It underscores the importance of considering maternal status as a key variable in studies of cognition and behavior. Furthermore, these findings have broader implications for understanding the neurobiological basis of cognitive flexibility and its potential modulation by hormonal and environmental factors. Future research could explore the molecular and cellular mechanisms underlying these cognitive changes and their relevance to human maternal behavior and clinical conditions characterized by cognitive impairments.

### Potential mechanisms how the reproductive status can affect cognition and motivation

Spatial learning and cognitive flexibility are complex cognitive functions regulated by interconnected neural circuits, including the hippocampus, and the prefrontal cortex. The hippocampus, in particular, is critical for spatial navigation and memory (Ekstrom & Hill 2023), while the prefrontal cortex supports executive functions such as cognitive flexibility and decision-making (Klune et al 2021). Both regions undergo significant plasticity during pregnancy and lactation, as evidenced by changes in dendritic structure, synaptic connectivity, and neurogenesis (Slattery & Hillerer 2016). However, the available evidence of neuronal plasticity in the postpartum period is more extensive in the hippocampus. Tomizawa et al. found that multiparous mice showed enhanced and sustained long-term potentiation (LTP) during the early postpartum period when compared to nulliparous mice (Tomizawa et al 2003). Cell proliferation and dendritic complexity are reduced in the hippocampus of maternal rats during the early postpartum phase, in contrast to virgin females (Darnaudery et al 2007). Additionally, postpartum neurogenesis has been linked to enhanced learning and memory in mothers. During weaning, when hippocampal neurogenesis is suppressed in rat dams (Pawluski et al 2006), spatial learning and memory show improvement (Pawluski et al 2006). Lactation itself may influence hippocampal neurogenesis, as the effects of pregnancy and birth on neurogenesis do not persist in the weeks following weaning (Medina & Workman 2020). Increasing nursing demand in rats led to reduced hippocampal neurogenesis, as evidenced by fewer immature neurons and decreased cell proliferation at weaning (De Guzman et al 2018). In addition to lactation, other factors may influence hippocampal neurogenesis, such as interaction with offspring. Nulliparous sensitized female rats have more neurogenesis in the hippocampus, both via cell proliferation and new cell survival, compared to either lactating rats or nulliparous female rats not exposed to pups (Pawluski & Galea 2007).

It is hypothesised that the alterations in neuroplasticity that occur during the postpartum period are attributable to the significant hormonal fluctuations experienced during pregnancy, labour, and the postpartum phase. The ensuing neuroplasticity may serve as the basis for the enhanced cognitive abilities observed in postpartum females. The postpartum-specific alterations in neuroplasticity are hypothesised to be driven by hormonal changes that are characteristic of this reproductive stage (Blankers et al 2024). Steroid and peptide hormones undergo significant fluctuations during the course of pregnancy, parturition and the postpartum period. Amongst these hormones, oxytocin is a potential mediator of enhanced hippocampus-dependent spatial learning in mothers. In a study by Tomizawa et al., oxytocin injections were found to enhance reference memory in nulliparous mice, yet no improvement was observed in working memory (Tomizawa et al 2003). Oxytocin, has also been suggested to play a role in improving cognitive flexibility because oxytocin receptor antagonism in the medial prefrontal cortex blocked the enhanced cognitive flexibility observed in mother rats (Leuner & Gould 2010). Additionally, ovarian hormones may also be involved, as high endogenous levels of estradiol (during proestrus) have been linked to impaired spatial learning, while low levels of estradiol have been associated with facilitated spatial learning (Holmes et al 2002). Estradiol levels increase as the pregnancy progresses. By late pregnancy, estradiol levels rise in both humans and rodents, and high estradiol is associated with worsened spatial abilities in female rats (Puri et al 2024). However, following parturition, estradiol levels decline and remain low during the lactational anoestrus period. This reduction in estradiol has been linked to enhanced performance in female rats (Henry & Sherwin 2012). With regard to the performance of working memory in mother mice, corticosterone may play a significant role. It has been demonstrated that females that have not been pregnant or given birth exhibit enhanced working memory performance when exposed to chronic stress, which is associated with elevated corticosterone levels (McLaughlin et al 2005). Given that basal corticosterone levels are elevated during lactation, it seems reasonable to posit that corticosterone plays an active role in spatial learning and memory enhancement. Another hormone with an elevated level in the postpartum period is prolactin. Surprisingly, the roles of prolactin, which is important for lactation and maternal behaviors, as well as adrenal hormones, in memory linked to the hippocampus during the postpartum phase, have not been fully explored (Moreno-Ruiz et al 2021).

In addition to hormones, it is also plausible that the environmental enrichment (pup exposure) contributes to the enhanced learning abilities of the mothers. However, it should be noted that the pups also serve as a source of stimulation for the control and pregnant females, given that they reside together in the IntelliCage with the mothers for an equivalent period. Consequently, a considerable proportion of the control females display sensitisation, exhibiting retrieval and licking of the pups despite not having given birth. Kinsley et al. observed that sensitised rats demonstrated enhanced performance on a reference memory task relative to nulliparous, but not primiparous rats (with mothering experience) following 16 days of pup exposure (Kinsley et al 1999). As compared to spatial tasks, executive functions like cognitive flexibility during pregnancy and motherhood in animal models have been less thoroughly studied. During the mid- and late postpartum periods, mothers show improved cognitive flexibility, a skill dependent on the prefrontal cortex. Executive functions were found enhanced during late pregnancy and the early postpartum period, but this improvement faded after weaning (Callaghan et al 2022). This effect required oxytocin signaling and the presence of the offspring. Blocking oxytocin receptors in the prefrontal cortex or removing the pups negates the positive impact of motherhood on cognitive flexibility. However, after weaning, motherhood no longer enhances performance on tasks involving set-shifting, even in rats that have gone through one or multiple pregnancies (Albin-Brooks et al 2017).

## Conclusions

In summary, the present study provides compelling evidence that the postpartum period in female mice is associated with pronounced enhancements in spatial learning, cognitive flexibility, and temporal anticipation, as demonstrated using the IntelliCage system. These findings build on previous observations by employing a long-term, minimally invasive, and ecologically valid monitoring method, offering novel insights into the cognitive adaptations of motherhood. The enhanced performance of the postpartum females across multiple tasks is indicative not only of heightened motivation but also of underlying neurobiological changes, likely driven by the interplay of hormonal fluctuations and environmental factors such as pup interaction. These enhancements, particularly in hippocampus- and prefrontal cortex-dependent functions, may confer evolutionary advantages by improving a mother’s ability to navigate her environment and respond effectively to the dynamic demands of caregiving. This research highlights the importance of investigating the neuroendocrine and experiential mechanisms that shape maternal brain plasticity, with potential implications for understanding human maternal behaviour and identifying novel targets for addressing cognitive impairments across clinical populations.

## Statements& Declarations

### Author Contributions

Melinda Cservenák: Conceptualization, Methodology, Validation, Formal analysis, Investigation, Data Curation, Writing - Original Draft, Writing - Review & Editing, Visualization; Luca Darai, Tamara Kállai, Bori Záhonyi: Methodology, Validation, Formal analysis, Investigation; László Détári: Data Curation, Writing - Review & Editing, Visualization; Arpád Dobolyi: Conceptualization, Methodology, Investigation, Resources, Writing - Original Draft, Writing - Review & Editing, Supervision, Project administration, Funding acquisition. All authors read and approved the final manuscript.

## Acknowledgments

We appreciate the technical assistance of Nikolett Hanák.

## Data Availability

The data presented in this study are available in the article, or are provided by the authors upon request.

## Ethics approval

This study was performed in line with the principles of the Declaration of Helsinki. Approval was granted by the Ethics Committee of Eötvös Loránd University (Date: 29.06.2021/No: PE/EA/568-7/2021).

## References

Agrati D, Lonstein JS. 2016. Affective changes during the postpartum period: Influences of genetic and experiential factors. Horm Behav 77: 141–52

Albin-Brooks C, Nealer C, Sabihi S, Haim A, Leuner B. 2017. The influence of offspring, parity, and oxytocin on cognitive flexibility during the postpartum period. Horm Behav 89: 130–36

Blankers SA, Go KA, Surtees DC, Splinter TFL, Galea LAM. 2024. Cognition and Neuroplasticity During Pregnancy and Postpartum In Neuroendocrine Regulation of Mammalian Pregnancy and Lactation, ed. PJ Brunton, DR Grattan, pp. 253-81. Cham: Springer International Publishing

Bramati G, Stauffer P, Nigri M, Wolfer DP, Amrein I. 2023. Environmental enrichment improves hippocampus-dependent spatial learning in female C57BL/6 mice in novel IntelliCage sweet reward-based behavioral tests. Front Behav Neurosci 17: 1256744

Callaghan B, McCormack C, Tottenham N, Monk C. 2022. Evidence for cognitive plasticity during pregnancy via enhanced learning and memory. Memory 30: 519–36

Cost KT, Lobell TD, Williams-Yee ZN, Henderson S, Dohanich G. 2014. The effects of pregnancy, lactation, and primiparity on object-in-place memory of female rats. Horm Behav 65: 32–9

Daguano Gastaldi V, Hindermann M, Wilke JBH, Ronnenberg A, Arinrad S, et al. 2025. A comprehensive and standardized pipeline for automated profiling of higher cognition in mice. Cell Rep Methods 5: 101011

Darnaudery M, Perez-Martin M, Del Favero F, Gomez-Roldan C, Garcia-Segura LM, Maccari S. 2007. Early motherhood in rats is associated with a modification of hippocampal function. Psychoneuroendocrinology 32: 803–12

De Guzman RM, Saulsbery AI, Workman JL. 2018. High nursing demand reduces depression-like behavior despite increasing glucocorticoid concentrations and reducing hippocampal neurogenesis in late postpartum rats. Behavioural Brain Research 353: 143–53

Diviney M, Fey D, Commins S. 2013. Hippocampal contribution to vector model hypothesis during cue-dependent navigation. Learning & memory 20: 367–78

Dobolyi A. 2025. Integrating the COM-B model into behavioral neuroscience: A framework for understanding animal behavior. Prog Neuropsychopharmacol Biol Psychiatry 138: 111346

Ekstrom AD, Hill PF. 2023. Spatial navigation and memory: A review of the similarities and differences relevant to brain models and age. Neuron 111: 1037–49

Gatewood JD, Morgan MD, Eaton M, McNamara IM, Stevens LF, et al. 2005. Motherhood mitigates aging-related decrements in learning and memory and positively affects brain aging in the rat. Brain Res Bull 66: 91–8

Habr SF, Bernardi MM, Conceição IM, Freitas TA, Felicio LF. 2011. Open field behavior and intra-nucleus accumbens dopamine release in vivo in virgin and lactating rats. Psychology & Neuroscience 4: 115–21

Henry JF, Sherwin BB. 2012. Hormones and cognitive functioning during late pregnancy and postpartum: a longitudinal study. Behav Neurosci 126: 73–85

Holmes MM, Wide JK, Galea LA. 2002. Low levels of estradiol facilitate, whereas high levels of estradiol impair, working memory performance on the radial arm maze. Behav Neurosci 116: 928–34

Izquierdo A, Brigman JL, Radke AK, Rudebeck PH, Holmes A. 2017. The neural basis of reversal learning: An updated perspective. Neuroscience 345: 12–26

Jennings JH, Ung RL, Resendez SL, Stamatakis AM, Taylor JG, et al. 2015. Visualizing hypothalamic network dynamics for appetitive and consummatory behaviors. Cell 160: 516–27

Johnson RF, Beltz TG, Thunhorst RL, Johnson AK. 2003. Investigations on the physiological controls of water and saline intake in C57BL/6 mice. Am J Physiol Regul Integr Comp Physiol 285: R394–403

Kinsley CH, Madonia L, Gifford GW, Tureski K, Griffin GR, et al. 1999. Motherhood improves learning and memory. Nature 402: 137–8

Kiryk A, Janusz A, Zglinicki B, Turkes E, Knapska E, et al. 2020. IntelliCage as a tool for measuring mouse behavior - 20 years perspective. Behav Brain Res 388: 112620

Klune CB, Jin B, DeNardo LA. 2021. Linking mPFC circuit maturation to the developmental regulation of emotional memory and cognitive flexibility. Elife 10

Lemaire V, Billard JM, Dutar P, George O, Piazza PV, et al. 2006. Motherhood-induced memory improvement persists across lifespan in rats but is abolished by a gestational stress. Eur J Neurosci 23: 3368–74

Leuner B, Gould E. 2010. Dendritic growth in medial prefrontal cortex and cognitive flexibility are enhanced during the postpartum period. J Neurosci 30: 13499–503

Lipp HP, Krackow S, Turkes E, Benner S, Endo T, Russig H. 2023. IntelliCage: the development and perspectives of a mouse- and user-friendly automated behavioral test system. Front Behav Neurosci 17: 1270538

Ma X, Schildknecht B, Steiner AC, Amrein I, Nigri M, et al. 2023. Refinement of IntelliCage protocols for complex cognitive tasks through replacement of drinking restrictions by incentive-disincentive paradigms. Front Behav Neurosci 17: 1232546

McLaughlin KJ, Baran SE, Wright RL, Conrad CD. 2005. Chronic stress enhances spatial memory in ovariectomized female rats despite CA3 dendritic retraction: possible involvement of CA1 neurons. Neuroscience 135: 1045–54

Medina J, Workman JL. 2020. Maternal experience and adult neurogenesis in mammals: Implications for maternal care, cognition, and mental health. J Neurosci Res 98: 1293–308

Moreno-Ruiz B, Mellado S, Zamora-Moratalla A, Albarracín AL, Martín ED. 2021. Increase in serum prolactin levels in females improves the performance of spatial learning by promoting changes in the circuital dynamics of the hippocampus. Psychoneuroendocrinology 124: 105048

Pawluski JL, Galea LA. 2007. Reproductive experience alters hippocampal neurogenesis during the postpartum period in the dam. Neuroscience 149: 53–67

Pawluski JL, Walker SK, Galea LA. 2006. Reproductive experience differentially affects spatial reference and working memory performance in the mother. Horm Behav 49: 143–9

Puri TA, Lieblich SE, Ibrahim M, Galea LAM. 2024. Pregnancy history and estradiol influence spatial memory, hippocampal plasticity, and inflammation in middle-aged rats. Horm Behav 165: 105616

Slattery DA, Hillerer KM. 2016. The maternal brain under stress: Consequences for adaptive peripartum plasticity and its potential functional implications. Front Neuroendocrinol 41: 114–28

Tomizawa K, Iga N, Lu YF, Moriwaki A, Matsushita M, et al. 2003. Oxytocin improves long-lasting spatial memory during motherhood through MAP kinase cascade. Nat Neurosci 6: 384–90

Uddin LQ. 2021. Cognitive and behavioural flexibility: neural mechanisms and clinical considerations. Nat Rev Neurosci 22: 167–79

Voikar V, Krackow S, Lipp HP, Rau A, Colacicco G, Wolfer DP. 2018. Automated dissection of permanent effects of hippocampal or prefrontal lesions on performance at spatial, working memory and circadian timing tasks of C57BL/6 mice in IntelliCage. Behav Brain Res 352: 8–22

